# Internal monitoring of whisking and locomotion in the superior colliculus

**DOI:** 10.1101/2024.03.16.585346

**Authors:** Suma Chinta, Scott R. Pluta

## Abstract

To localize objects using active touch, our brain must merge its map of the body surface with an ongoing representation of self-motion. While such computations are often ascribed to the cerebral cortex, we examined the midbrain superior colliculus (SC), due to its close relationship with the sensory periphery as well as higher, motor-related brain regions. We discovered that active whisking kinematics and locomotion speed accurately predict the firing rate of mouse SC neurons. Kinematic features occurring either in the past, present, or future best predicted spiking, indicating that the SC population continuously estimates the trajectory of self-motion. Half of all self-motion encoding neurons displayed a touch response as an object entered the active whisking field. Trial-to-trial variation in the size of this response was explained by the position of the whisker upon touch. Taken together, these data indicate that SC neurons linearly combine an internal estimate of self-motion with external stimulation to enable active tactile localization.

## Introduction

In the absence of body movement, tactile localization is performed by mapping stimuli directly to receptors on the body surface. The somatosensory system of rodents contains an elegant map of receptor space, where each whisker on the face activates a corresponding coordinate in the brain (*1–7*). However, in real life, the position of the body constantly changes as animals explore their environment and orient towards stimuli (*8, 9*). Therefore, active tactile localization requires the brain to combine its somatotopic map of the whiskers with an egocentric model of their position (*10*). Where in the brain does this joint representation of somatotopic space and whisker position arise?

The midbrain superior colliculus (SC) contains a somatotopic map of the whiskers and is hypothesized to play an essential role in whisker-guided movement (*11–14*). It receives monosynaptic input from the vibrissae regions of the trigeminal nucleus, somatosensory and motor cortices, as well as the cerebellum, making it an ideal candidate for combining efferent and afferent signals (*15–19*). Whisker movement is encoded in the somatosensory system through reafference: the self-generated activation of receptors in the follicle (*20*). During rhythmic whisking, neurons in the trigeminal ganglion, brainstem, thalamus, and cortex encode the relative position of the whisker within a cycle of movement, termed phase (*21–25*). Neurons in the SC are known to spike at the onset of electrically induced (fictive) whisking, yet their functional relationship to phase is unknown (*26*).

While phase information is necessary for encoding the rhythmic changes in position that occur within a whisk cycle, midpoint and amplitude information are necessary to build a complete model of whisker position (*10, 12, 27*). SC neurons could conceivably use or build such model, since they receive input from whisker phase (barrel cortex and brainstem), amplitude and midpoint (motor cortex and cerebellum) encoding brain areas (*18, 22, 27–34*). Despite these known anatomical connections, the relationship between SC spiking and volitional whisker position is unknown.

The SC is hypothesized to play a key role in generating whisker movement, since a sparse population of its excitatory neurons target the downstream nuclei that control the whisking muscles (*35–37*). Functional evidence supporting a role for the SC in whisker movement comes from its electrical activation under anesthesia, which causes sustained whisker protraction (*38, 39*). Likewise, unilateral ablation of the SC retracts the resting position of the contralateral whisker pad but has no effect on volitional bouts of whisking (*35*). These data argue that in the absence of whisking, SC spiking is directly related to whisker midpoint. However, recent evidence reveals that SC neurons also have excitatory connections with the nucleus that mediates whisker retraction (*35, 37, 40*). Therefore, the actual role of the SC in whisking remains unclear. Moreover, the tactile features encoded by movement-related SC neurons are unknown. Given the established role of the SC in visuomotor processing (*41–45*), we hypothesized that neurons in the mouse SC combine an internal representation of whisker movement with tactile information to enable active tactile localization (*10, 12, 46*).

To determine the kinematic and tactile features encoded by SC neurons, we performed high-density electrophysiology and high-speed videography in head-fixed, freely whisking and locomoting mice that were periodically presented with an object surface. We discovered that the firing rates of many SC neurons were predicted by the kinematics of whisking and locomotion. SC activity was best predicted by movements occurring either in the present or future time domains, indicating that the SC is important for the adaptive control of movement. Neural selectivity for a combination of kinematic features enabled an accurate representation of absolute whisker angle. Half of all neurons encoding self-motion also displayed a significant tactile response. Trial-to-trial variation in the size of this response was correlated to the position of the whisker upon touch. Taken together, these data indicate that SC neurons combine an internal estimate of body position with external stimulation to enable active tactile localization.

## Results

### The firing rates of SC neurons are linearly related to self-motion

To reveal the neural representation of whisking and locomotion, we recorded single-unit activity from populations of neurons located in the deep layers of the lateral SC (Fig. 1a and Suppl. Fig. 1a). Mice were head-fixed yet able to whisk and locomote freely on a treadmill, allowing for behaviors that were under complete volitional control (*47–50*). In an average recording session, we imaged whisking for 600 ± 167 seconds, where mice completed 5857 ± 1507 whisk cycles and locomoted 103 ± 28 meters. Using high-speed (500 fps) infrared imaging and markerless tracking (*51*), we calculated whisker position over time, and decomposed its movement into distinct kinematic features, known as midpoint, amplitude, and phase (Fig. 1b, see Methods). We categorized these whisking features as ‘slow’ or ‘fast’, based on the derivative of their autocorrelations (Suppl. Fig. 1b), and deemed whisker midpoint and amplitude as slow, relative to the much faster (<1 whisk cycle) movements that underlie phase (Fig.1c).

**Figure 1.**
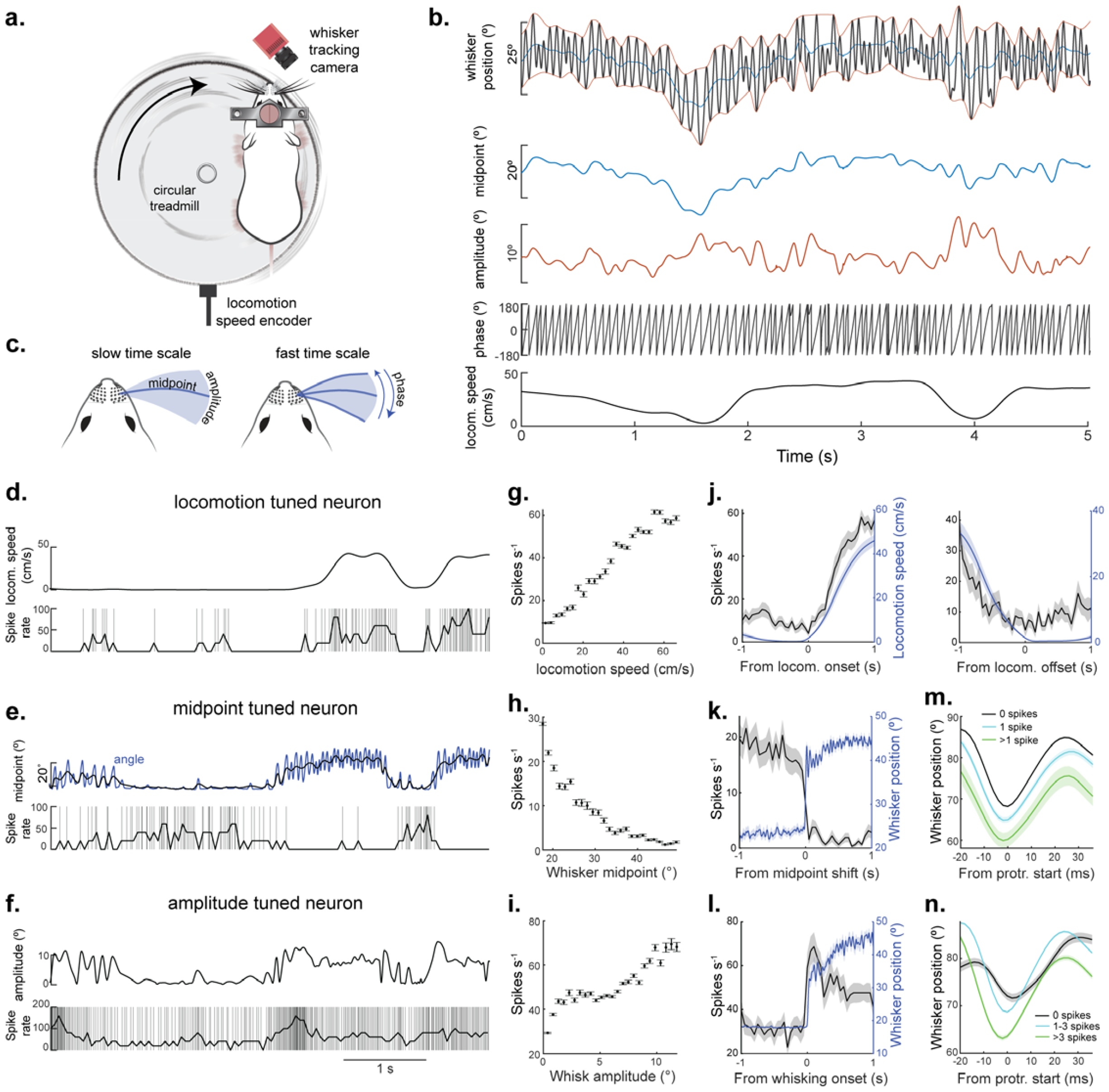
The firing rate of SC neurons is linearly related to self-motion. **A**) A schematic of the experimental setup illustrating a head-fixed mouse locomoting on a circular treadmill and actively whisking in air. A high-speed camera and digital encoder recorded whisker position and locomotion speed, respectively. **B**) Example traces of whisking kinematics and locomotion speed over time. Whisking features midpoint, amplitude, and phase were calculated from angle. **C**) Illustration displaying the slow-changing features whisker midpoint and amplitude (left) and the fast-changing feature whisker phase (right). **D – F**) The relationship between self-motion (locomotion (**d**), midpoint (**e**) or amplitude (**f**)) and spike rate in three example neurons. Each row of plots corresponds to a different neuron. **G**) Firing rate as a function of locomotion speed for the neuron in **d. H**) Firing rate as a function of whisker midpoint for the neuron in **e. I**) Firing rate as a function of whisk amplitude for the neuron in **f. J**) Neuronal firing rate aligned to locomotion onset and offset (left and right) for the neuron in **d. K**) Whisker position (blue) and firing rate (black) aligned to when the mouse made a large increase in midpoint. **L**) Firing rate (black) aligned to the onset of whisking (blue) for the neuron in **f. M**) Mean whisker position (aligned to protraction start) preceded by either 0, 1, or >1 spike in the neuron in **e. N**) Mean whisker position (aligned to protraction start) preceded by either 0, 1 – 3, or >3 spikes in the neuron in **f**.

To determine the relationship between SC spiking and the slow kinematic features (including locomotion), we first used a traditional tuning curve approach. In many neurons, we observed a linear relationship between firing rate and the kinematics of whisking and locomotion speed (Fig, 1d – i & Video 1). To better understand the temporal fidelity of this relationship, we plotted firing rate relative to transitions in self-motion (locomotion onset/offset, midpoint shift, whisking onset), revealing precise and accurate coupling between spike timing and behavior (Fig. 1j – l). Furthermore, we segregated whisk cycles according to the number of spikes in the preceding whisk cycle (Fig. 1m, n) and found that the number of preceding spikes was correlated to the midpoint/amplitude in the upcoming whisk. Taken together, these data highlight the strong and temporally precise correlation between the activity of SC neurons and self-motion. While this analysis provides an intuitive basis for understanding the relationship between neural activity and self-motion, it fails to capture nonlinear dynamics or address the inherent correlation between kinematic features. To address these issues and objectively quantify the dependence of SC spiking on the different features of self-motion, we implemented a previously established linear-nonlinear Poisson (LNP) spiking GLM (*52, 53*). Given the natural correlation between kinematic features, the LNP model with forward search is advantageous because it identifies the kinematic feature(s) that provide(s) a significant contribution to spike prediction (Suppl. Fig. 2) (*50, 54*).

**Figure 2.**
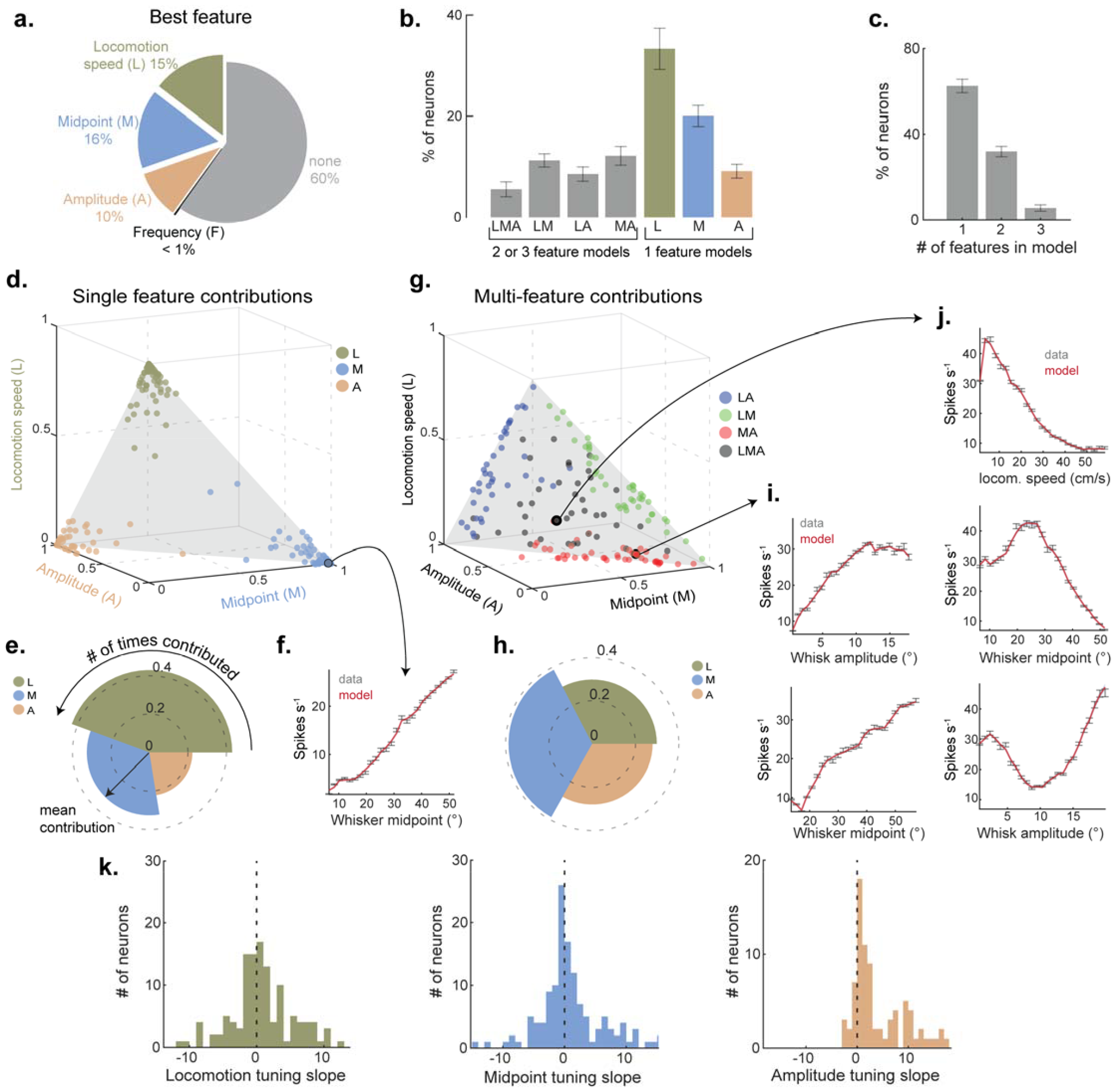
Single neuron encoding of whisking and locomotion. **A**) The kinematic feature that best predicted spiking across the recorded population (feature with largest log-likelihood among single feature models). **B**) Percentage of neurons with single and multifeature selectivity for whisking and locomotion (352/932 neurons, 12 mice). **C**) Percentage of neurons selective to 1, 2, or 3 kinematic features (352 neurons). **D**) Scatterplot of normalized contributions of individual features (locomotion speed, midpoint, and amplitude) to selected model of single-feature selective neurons (187 neurons). **E**) Polar plot denoting the relative contribution of individual features to the full model for single-feature selective neurons (187 neurons). **F**) Neuronal tuning curve to midpoint (best feature selected by model). **G**) Scatterplot of normalized contributions of individual features to selected model of multi-feature selective neurons (165 neurons). **H**) Polar plot denoting the relative contribution of individual features to the full model for multi-feature neurons (165 neurons). **I, J**) Tuning curves of two neurons with significant contributions from multiple kinematic features. **K**) Histogram of neuronal tuning curve slopes to different self-motion features (p=0.01, 125 neurons whose best feature was locomotion speed; p=0.01, 136 neurons whose best feature was midpoint; p=1.5e^-9^, 83 neurons whose best feature was amplitude; one-sample t-test).

### Mixed selectivity for whisking and locomotion

To identify the self-motion features that predict SC spiking, we fit every neuron with single- and multi-feature LNP models. Using the forward search approach, we identified the model with the least number of features that best predicted spiking. Any kinematic feature that did not significantly boost model performance over the best single-feature model was discarded (Suppl. Fig. 2c – e). This approach revealed that 40% of SC neurons had a firing rate predicted by one or more slow self-motion features (345/859 neurons in 12 mice, Fig. 2a, b). The activity of most neurons was best predicted by whisker midpoint or locomotion speed (16% and 15% of neurons, respectively), while a smaller subset of neurons preferred whisk amplitude (10%). For most neurons tuned to self-motion (59%, 180/345), only a single kinematic feature provided a significant contribution to spike prediction, while the remaining neurons benefited from a combination of features (41%, 165/345, Fig. 2b, c).

Amongst the single-feature neurons, locomotion speed predicted spiking most often (Fig. 2d, e). However, amongst neurons with multi-feature tuning, whisker midpoint emerged as the most frequent and strongest contributor to spike prediction (Fig. 2g, h). These multi-feature neurons could be essential for building an accurate model of whisker position (as in midpoint + amplitude neurons: MA) or performing behaviors that rely on simultaneous adjustments to whisking and locomotion (as in locomotion + X neurons: LM, LA, or LMA).

Across the population, the tuning slopes for locomotion speed and whisker midpoint exhibited a bidirectional distribution (Fig. 2k), suggesting a nuanced role for the SC in movement control, where increases in spike rate could either facilitate (positive slopes) or suppress (negative slopes) movement. This may enable the SC to delicately balance the initiation and cessation of actions while controlling appetitive motion (*41, 55– 58*). Spike rate was positively correlated to whisk amplitude, indicating that SC activity could have an important role in adapting the whisking strategy between large exploratory and small foveal movements (*59*).

### Past, present, and future movements predict SC spiking

Active sensing is a recurrent process that integrates knowledge of the body’s current state with future predictions (*8, 60, 61*). Such computations are critical for adapting movement trajectories based on environmental cues (*9, 11, 47, 62*). To determine if the SC is capable of this computation, we utilized time-shifted models and tested the preference of SC neurons for past, present, and future kinematic features. SC neurons displayed one of three temporal preferences. Units biased toward the past exhibited relatively steady model performance for past time shifts, contrasted with a rapid decay in performance for the future (Fig. 3a). In some neurons, peak model performance also occurred in the past, underscoring their prominent role in storing traces of prior motion. Present biased units showed a symmetric decline in model performance, reflecting a real-time representation of self-motion (Fig. 3b). In these units, spike timing suggests a role either in immediate sensory feedback or an efferent copy of motor commands. Future biased units displayed steady model performance for future time shifts but a rapid decay for the past (Fig. 3c). This pattern implies a predictive model of future body position. Unsupervised (k-means) clustering of neuron response profiles confirmed these categories, distinguishing three principal clusters corresponding to past, present, and future biased populations (Fig. 3d). The distribution of temporal biases and preferred time shifts indicate that the SC population represents a trajectory of body position spanning up to several hundred milliseconds (Fig. 3e, f, g). The temporal bias of neurons was similar between the different kinematic features (Suppl. Fig. 3). Overall, these data reveal a preference for present and future time domains. A large fraction of SC neurons preferred time-shifts within 50ms of the present, a timescale relevant to whisking (mean whisk period is 52ms). These data support a role for the SC in adaptive motor of whisking (*41, 63–65*).

**Figure 3.**
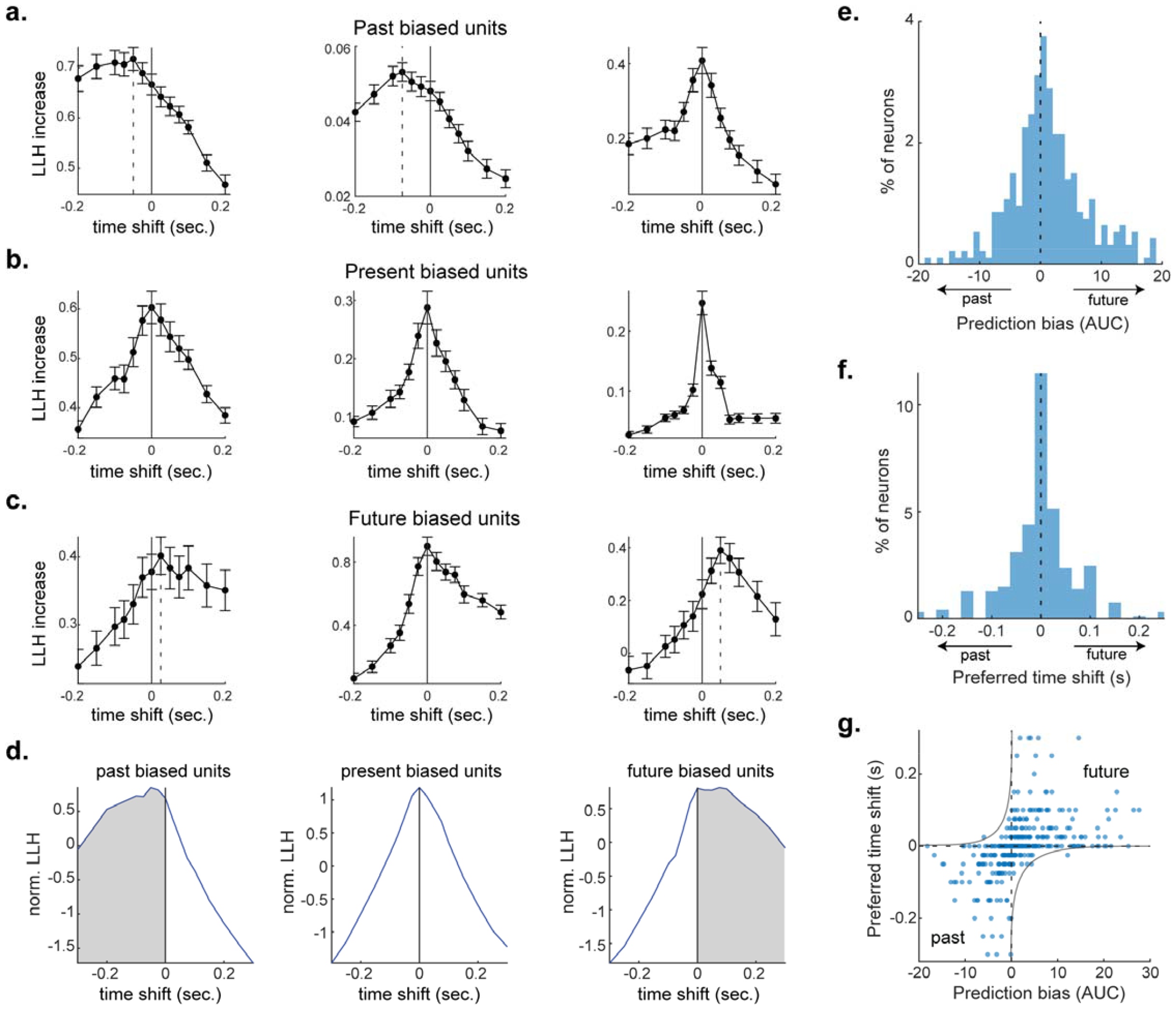
The SC encodes past, present, and future movements. **A**) Three example neurons with firing rates more accurately predicted by past movement features. The y-axis measures the log-likelihood (LLH) of spike prediction, which reflects the accuracy of the neural encoding model. The x-axis displays the time shift of the self-motion feature relative to the onset of neural activity, ranging from -0.2 seconds (indicative of the past) to +0.2 seconds (indicative of the future). **B, C**) Example neurons with spiking most accurately predicted by present (**B**) or future (**C**) self-motion features. All error bars represent mean +/-sem. **D**) Unsupervised clustering of time-shifted neuronal encoding curves created three clusters that segregate into past, present, and future biased units (78, 123, and 104 neurons respectively). The templates of these clusters are displayed. **E**) Distribution of temporal biases which measures the area under the curve for future times relative to past times (12 mice, 347 neurons). **F**) Distribution of preferred time shifts (12 mice, 347 neurons). **G**) Relationship between temporal bias and preferred time shift (12 mice, 347 neurons).

### The SC contains a comprehensive map of whisker phase

In exploring the representation of self-motion, we extended our analysis beyond the slow-changing features to investigate neural selectivity to the phase of whisker motion. Phase tuning has been observed throughout the ascending pathway of the somatosensory whisker system, but it has never before been investigated in the SC (*21, 23–25*). To examine phase tuning in SC neurons, we aligned spikes to the onset of whisker protraction, and we also generated spike triggered averages of phase angle using an established approach (*22, 23*) (Fig. 4c, d). Many neurons displayed non-uniform firing rates, often having a clear preference for either the protraction or retraction period of the whisk cycle. Across the population, 38% of neurons were significantly tuned to phase (327/859, chi-squared test, p < 0.05), with the most frequent phase preference occurring immediately after the start of retraction (Fig. 4e & f). Nonetheless, the overall distribution of neuronal preferences covered the entire range of phase angles (Fig. 4g). Half of all neurons tuned to phase also encoded one or more slow self-motion features (47%, 161/345). This is exemplified in Figure 4h & i, by showcasing a neuron with phase tuning modulated by midpoint and amplitude. We demonstrate this slow-feature scaling of the phase response by evenly dividing phase angles between their upper and lower percentiles of midpoint or amplitude (Fig. 4j). This fast + slow coding scheme theoretically supports the computation of absolute whisker angle. We experimentally tested this theory below using a standard decoder model.

**Figure 4.**
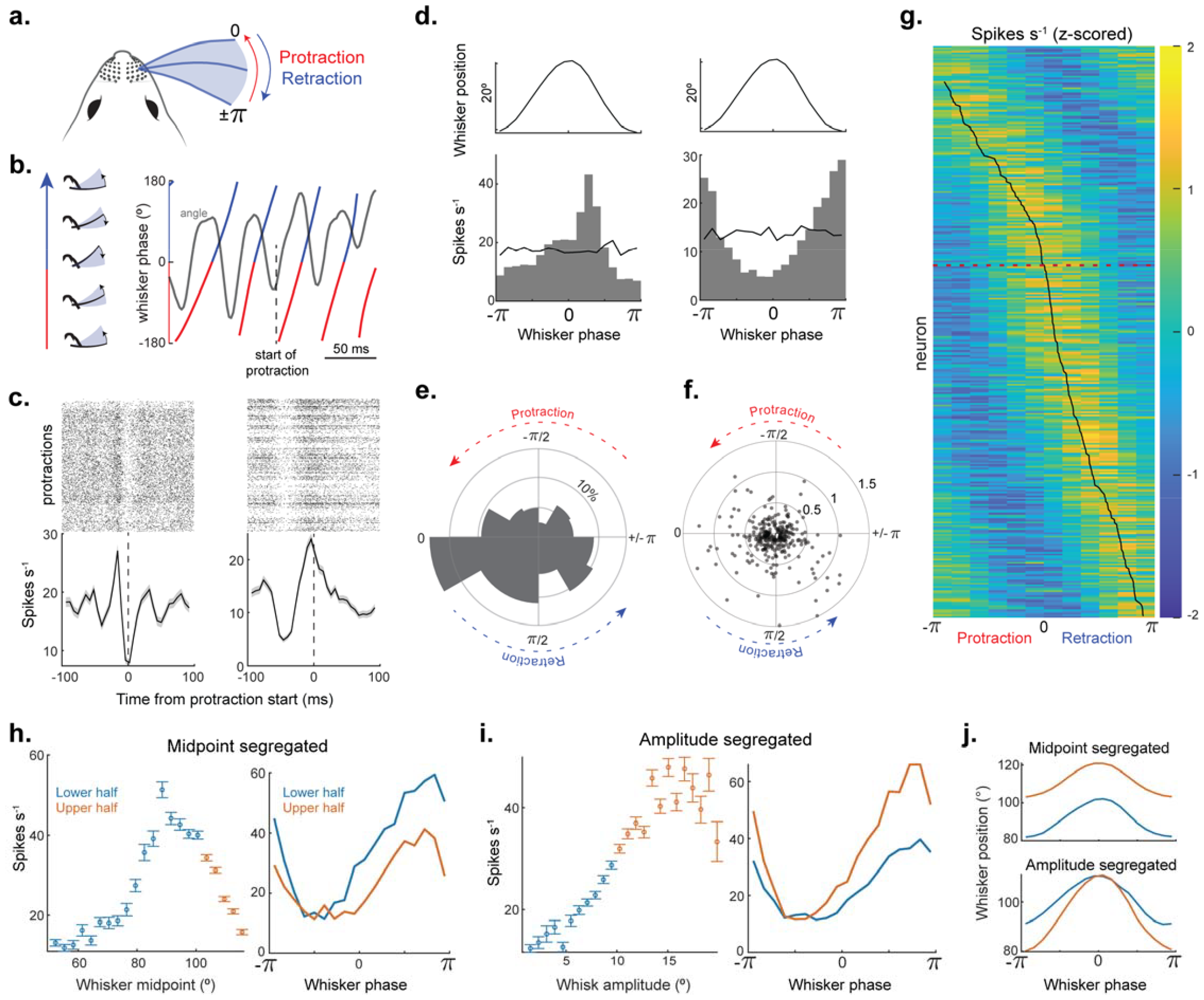
SC neurons encode the phase of whisker motion in combination with slow self-motion features. **A**) Schematic of relationship between whisker position and phase. Protraction period of the whisk cycle is represented in red and the retraction period in blue. **B**) Aligned traces of whisker position (in grey) and whisker phase (color). Dashed line represents the start of protraction in a whisk cycle. **C**) Spike raster (top) and mean firing rate (bottom) aligned to the start of whisker protraction for two example neurons. **D**) Top: whisker position as a function of phase. Bottom: phase tuning curves for the two neurons in panel **C**. The solid black line indicates the neuron’s firing rate if spike times were randomly distributed. **E)** Polar histogram of preferred phase angle across the population of significantly tuned neurons (p<0.05, 12 mice, 327 neurons, chi-squared test). **F**) Preferred phase and modulation depth for each neuron (327 neurons). **G**) Heat map of phase tuning curves aligned by their preferred phase (12 mice, 327 neurons). **H**) Left: Whisker midpoint tuning for an example neuron that encodes whisker midpoint, amplitude, and phase. Midpoints are segregated into upper (orange) and lower (blue) halves. Right: Whisker phase tuning of the neuron in **H** segregated by upper and lower midpoints. **I**) Left: Whisker amplitude tuning for neuron in **H** color-coded for upper and lower halves. Right: Whisker phase tuning of the neuron in **H** segregated by upper and lower whisk amplitudes. **J**) Whisker position as a function of phase for midpoint (top) and amplitude segregated (bottom) whisk cycles.

### The SC computes the absolute angle of whisker position

We implemented a decoder model to predict absolute whisker angle from SC population activity spanning past, present, and future time bins (Fig. 5a, see methods). For each 175 ms period of SC activity, whisker position was predicted in a 15 ms window, to allow for the capture of rapid, sub-cycle changes in whisker position.

**Figure 5.**
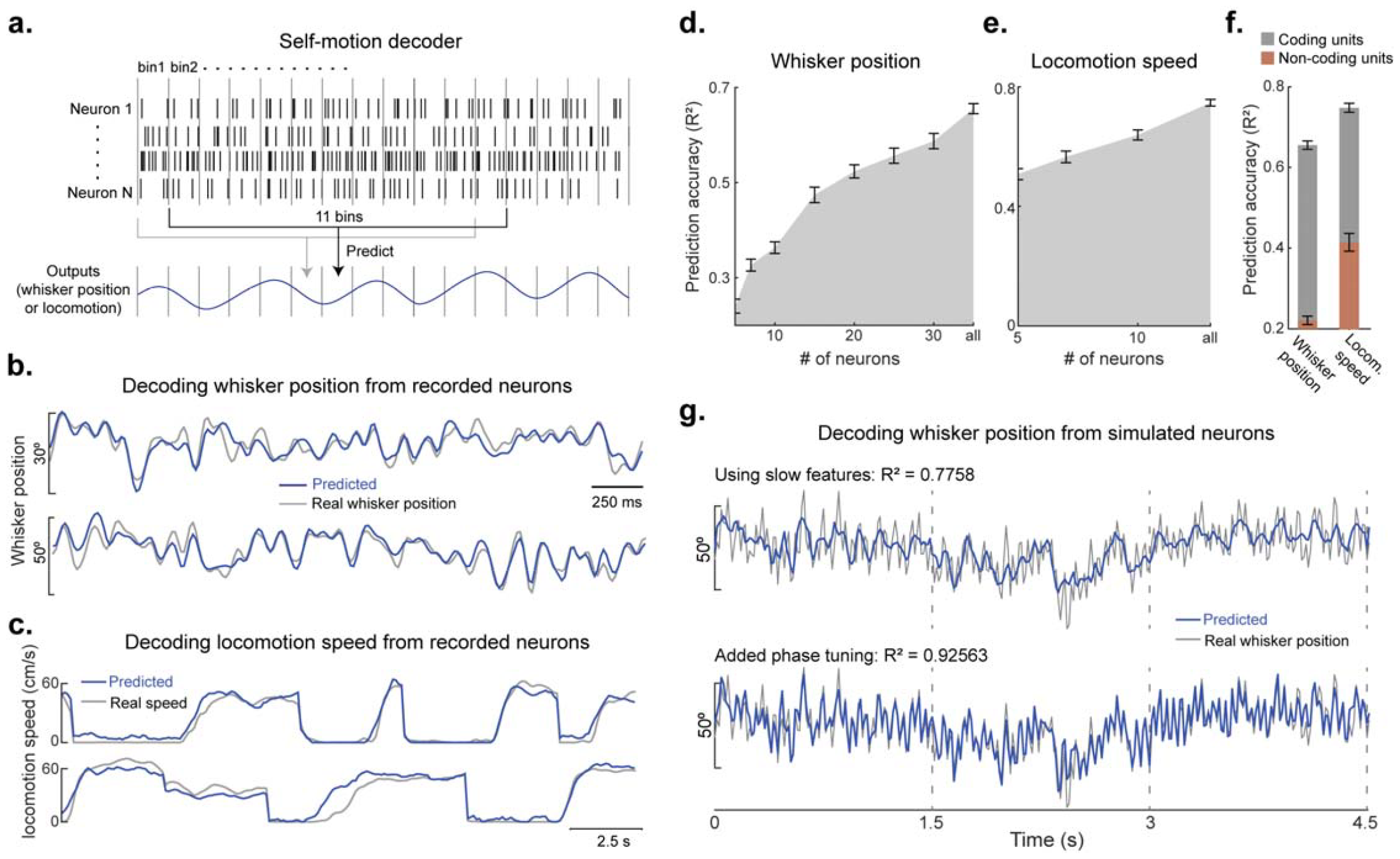
SC spiking accurately predicts whisker angle and locomotion speed. **A**) Schematic of self-motion decoder. To predict the self-motion feature in each time bin (15 ms for whisker position and 100 ms for locomotion speed), the firing rate of N neurons in T time bins were used (T=11 for whisker position, 9 for locomotion speed). **B**) Prediction of whisker position from all whisking selective neurons (as determined by LNP model) in a recording session (49 neurons, mouse 1; 35 neurons, mouse 2). **C**) Prediction of locomotion speed from all locomotion selective neurons in a recording session (16 neurons, mouse 1; 11 neurons, mouse 2). **D, E**) Prediction accuracy of whisker position (in **D**) and locomotion speed (in **E**) as a function of increasing number of neurons in the decoder model (7 mice in D, and 10 mice in E). Only neurons with significant self-motion tuning were used. **F**) Prediction accuracy of whisker position and locomotion speed for self-motion coding and non-coding units (12 mice). **G**) Top, decoding whisker position from simulated neurons that only encode slow self-motion features (n = 40 neurons). Bottom, decoding whisker position after the addition of simulated neurons that are tuned to whisker phase (bottom) (n = 40 neurons, 20 encode slow self-motion, 20 encode phase tuning).

Remarkably, whisker position was predicted with a high level of accuracy, as evidenced in snapshots from two example mice (Fig. 5b) and overall model performance across the population of mice (Fig. 5d and Suppl. Fig. 4). To our knowledge, the accuracy of this result surpasses other tested brain regions. However, given the uniquely large size of our dataset, encompassing several thousand whisk cycles in each mouse, such comparisons are difficult to interpret. Interestingly, SC activity was also adept at decoding locomotion speed (Fig.5c). Decoder performance was robust, even with a modestly sized population of units, and performance gradually increased with the number of neurons included in the model (Fig. 5d, e). Neurons in our recorded population that did not encode self-motion (as determined by the LNP model) provided very little predictive power for whisker position (Fig. 5f, orange bars). To disentangle the impact of slow and fast (phase) kinematic tuning on decoding whisker position, we simulated the activity of multiple neurons that closely mirrored our experimental observations (see methods). Results from this simulation revealed that neurons tuned exclusively to slow self-motion features (midpoint and amplitude) can reliably predict gradual changes in whisker position. Next, by incorporating neurons with phase tuning, model performance was markedly enhanced, by capturing the rapid, sub-cycle changes in whisker position (Fig. 5g, R^2^ = 0.92 vs. R^2^ = -0.1 for shuffled spikes in Suppl. Fig. 4e). Taken together, these data illustrate the importance of phase tuning and the complementary nature of slow and fast self-motion tuning for generating a high-resolution map of absolute whisker angle in the SC.

### The representation of self-motion is modified by sensory reafference

To determine if sensory reafference plays a significant role in the representation of self-motion, we trimmed the length of the whiskers (to less than 4 mm), significantly diminishing the inertial forces on the follicle (*21*) (Fig. 6a). To examine the impact of trimming on different kinematic features, we calculated LNP model performance pre- and post-trimming. To control for changes in behavior between the trimming conditions, we only compared model performance across equivalent feature space. A notable portion of neurons maintained similar model performance pre- and post-trimming, suggesting a minor role for sensory reafference in these neurons (Fig. 6b & g, 36% of neurons for slow features). Conversely, other neurons exhibited a significant change in model performance (increase or decrease), and some neurons lost or gained the ability to encode self-motion (Fig. 6c, d, e, & f). To test the influence of re-afference on phase tuning, we calculated the Pearson correlation between the pre- and post-trimming tuning curves of each neuron (Fig. 6h). Comparable changes in tuning were observed for phase (Fig. 6a – e bottom row and f). Overall, trimming caused a significant reduction in firing rates, even when controlling for whisker position and locomotion speed occupancies between the conditions (Fig. 6i, 8 mice, 636 neurons, Wilcoxon signed rank test, p = 2e-26). These data reveal that sensory reafference shapes the representation of self-motion in SC neurons.

**Figure 6.**
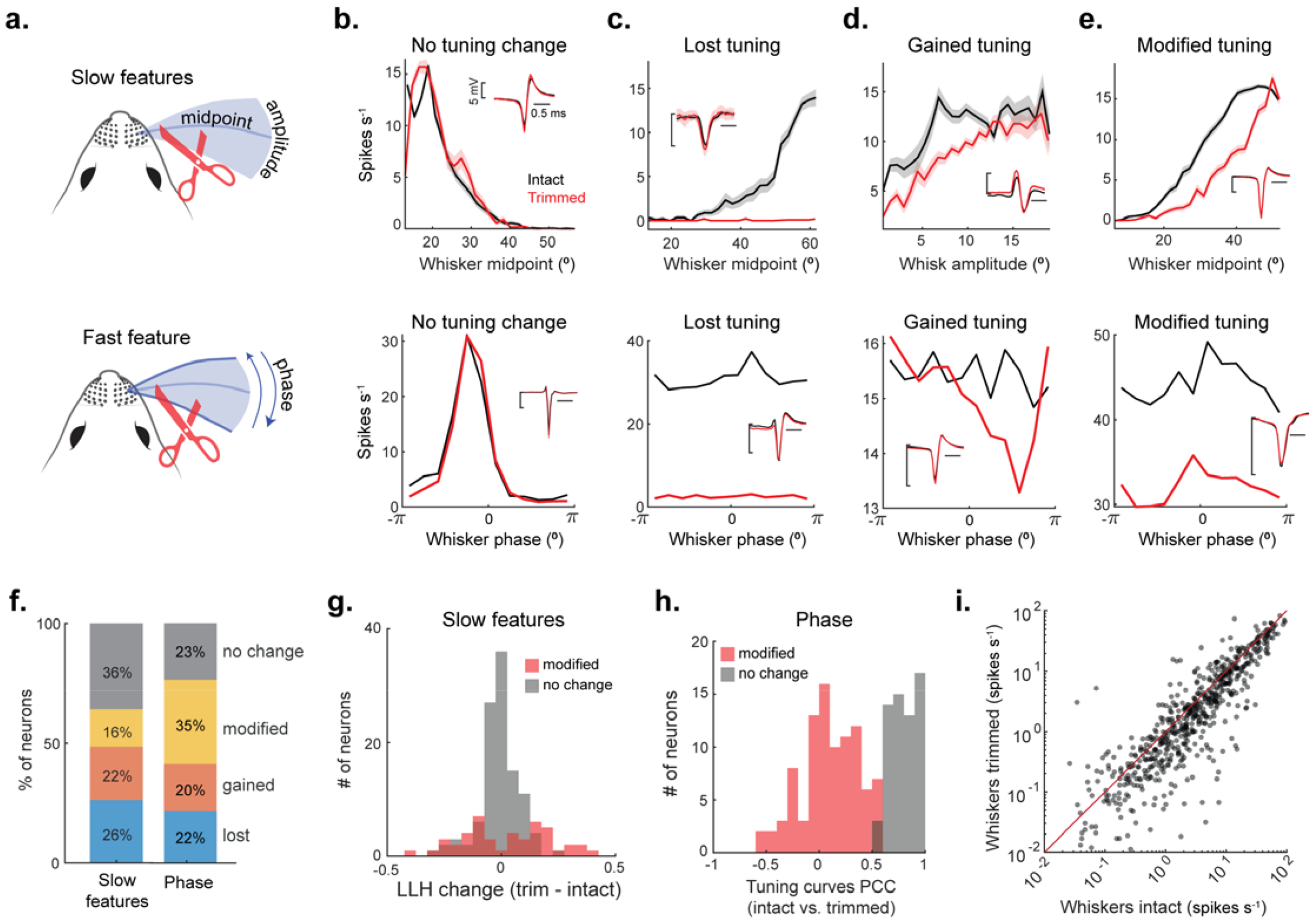
The representation of self-motion is modified by sensory reafference. **A**) A schematic showing whisker trimming and self-motion features. **B – E**) Top: four example tuning curves for slow self-motion features during free whisking (black) and after trimming off the whiskers (red). Insets are spike waveforms during the two conditions. Bottom: four example tuning curves to whisker phase during free whisking (black) and after trimming (red). **F**) Percent of units that lost, gained, modified, or had no change in slow-or fast-feature tuning after whisker trimming (slow: 8 mice, 264 neurons; fast: 8 mice, 301 neurons). **G**) Change in log-likelihood (LLH) values of each neuron after whisker trimming. Units in red have significantly modified LLH after trimming (one-tailed t-test, p<0.05, 47 neurons; no change, p>0.05, 108 neurons). **H**) Pearson correlation coefficient (PCC) between phase tuning curves calculated before and after whisker trimming. Units in red have significantly modified phase tuning curves (PCC, p<0.05, 62 neurons; no change, p>0.05, 93 neurons). **I**) Firing rates of SC neurons during whisking and locomotion calculated before and after whisker trimming. The whisker position and locomotion speed occupancies were selected to be equivalent between the conditions (8 mice, 636 neurons, Wilcoxon signed rank test, p = 2e-26).

### The integration of internal and external representations

To compute the egocentric distance of an object during active touch, the brain must combine tactile information with an internal estimate of body position. To investigate this computation in SC neurons, we examined the tactile response as an object moved into (12.6 cm/s) the active whisking field (Fig. 7a & Suppl. Video 1). Using high-speed imaging, we calculated the position of the whisker during the transient tactile response (Fig. 7b).

**Figure 7.**
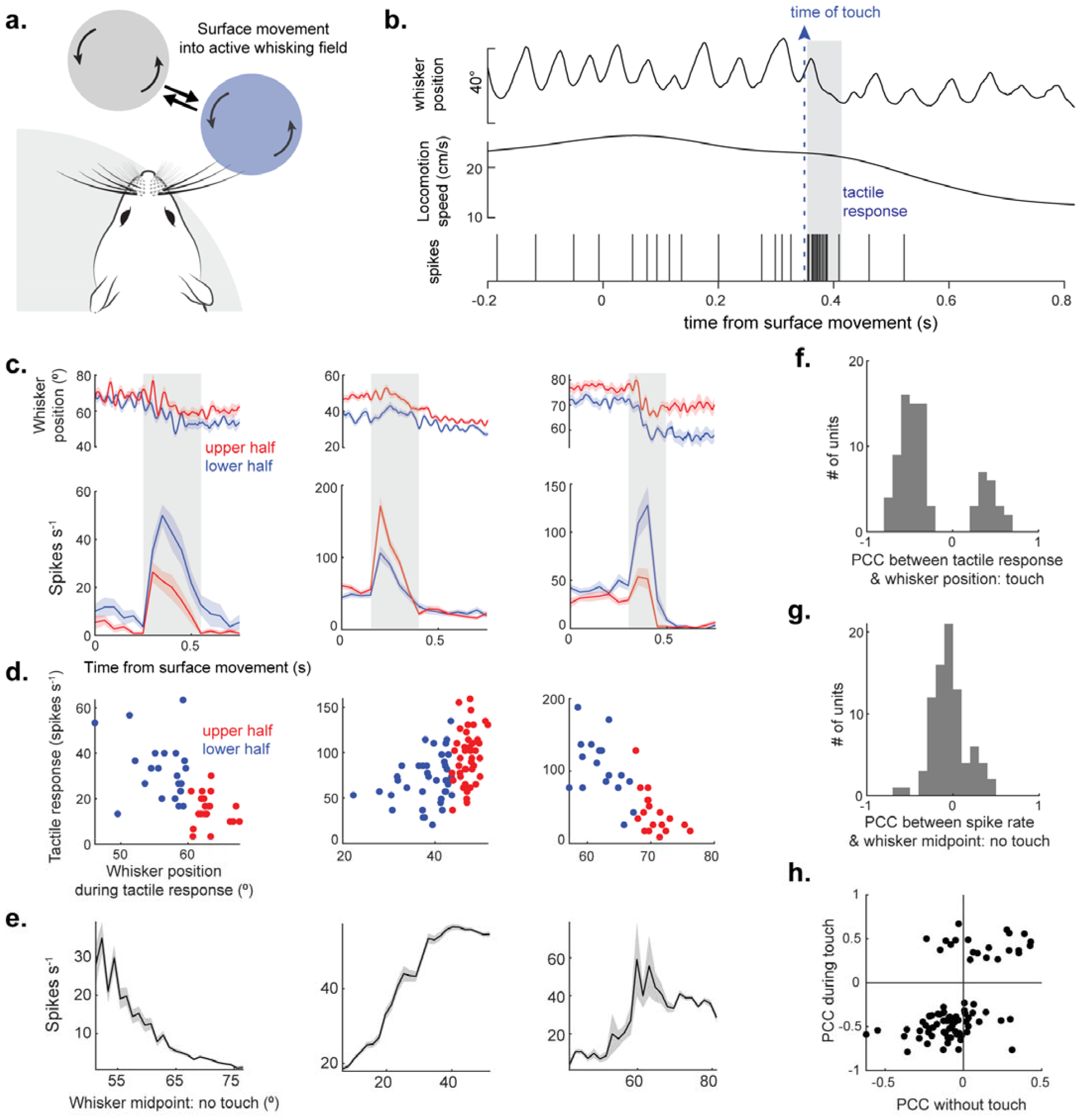
Whisker position directly correlates with tactile response gain. **A**) A schematic illustrating object movement into the active whisking field of the mouse. **B**) Whisker position, locomotion speed and spiking in an example neuron as the object moves into the whisker field. Dashed blue line indicates the onset of whisker touch with the surface. The gray window indicates the neuron’s response period. **C**) Three example neurons with a strong correlation between the tactile response magnitude and whisker position. Whisker position (top) and firing rate (bottom) during touch. The tactile response window is marked in gray. The upper half of whisker positions trials are marked in red, and the lower half are in blue. **D**) Scatter plot of mean spike rate vs mean whisker position for the three neurons in **C**, calculated on a trial-by-trial basis (Pearson correlation coefficient = -0.61,0.46, and -0.76 respectively). **E**) Free-whisking midpoint tuning curves for neurons in **C** (Pearson correlation coefficient = -0.38, 0.43, and 0.31 respectively). **F**) Distribution of significant (p < 0.05) Pearson correlation coefficients (PCCs) between spike rate and whisker position (12 mice, 83 neurons). **G**) Distribution of PCCs between spike rate and whisker position during free-whisking for neurons in **F** (12 mice, 83 units). **H**) Scatter plot of the correlation coefficient between spike rate and whisker position, calculated during the tactile response period (y-axis) and during free whisking (x-axis, 83 neurons).

The transient response period was manually selected for each mouse by identifying the period when the population-level activity was greater than baseline. First, we divided firing rates into two halves (upper vs. lower), determined by the mean position of the whisker during the tactile response, as demonstrated in three example neurons (Fig. 7c). We discovered that whisker position upon touch (upper vs. lower half) was correlated to tactile response magnitude, with either a positive or negative relationship. Next, we correlated the tactile response of these neurons with whisker position on a trial-by-trial basis and discovered a strong, linear trend (Fig. 7d). Interestingly, tuning to whisker midpoint (calculated during free whisking) in these neurons followed a similar trend (Fig. 7e). Overall, 20% of neurons with a tactile response displayed a significant linear correlation between response magnitude and whisker position (Pearson correlation, p < 0.05, Fig. 7f, n = 83). The sign of this effect (positive or negative) often mirrored the correlation we observed during free whisking (Fig. 7g & h). In these neurons (n = 83), neither whisker curvature nor habituation better explained the dependence of the tactile response on whisker position (Suppl. Fig. 5). Neurons that showed a stronger relationship with trial repetition (habituating neurons), as shown in our previous study (*49*), were removed from this analysis. These data indicate that the ongoing internal representation of self-motion directly modulates the strength of the tactile response.

## Discussion

We discovered that the kinematics of active whisking and locomotion accurately predict the spiking of SC neurons. Our analysis revealed populations of neurons that were maximally responsive to movements either in the past, present, or future. Therefore, the SC creates a continuous sensorimotor representation of movement trajectories. This internal monitoring of whisking operated across timescales, with half of all self-motion neurons encoding a combination of slow and fast kinematic features. This comprehensive map of self-motion accurately predicted absolute whisker angle. Sensory reafference played a key role in building this model. Lastly, tactile response magnitude was linearly related to the position of the whisker upon active touch. This interaction between internal and external representations could enable SC neurons to compute the egocentric distance of an object during touch.

Many studies have demonstrated that locomotion modulates neural activity, but how this modulation changes as a function of speed has often been overlooked (*66–74*). Our dataset, collected across a wide range of locomotor speeds, revealed a diversity of tuning profiles. SC firing rates have been known to increase at the onset of locomotion (*57*), but we found a broad range of positive and negative tuning curves that could predict locomotion speed with incredible accuracy. Interestingly, many SC neurons simultaneously encoded whisking and locomotion (LNP model performance significantly improved with both features). This multi-feature representation is likely useful for controlling appetitive movements and may bestow animals with a more seamless control over sensory-guided locomotor decisions (*50, 54, 58, 75–78*). Non-coding SC neurons poorly predicted (R^2^ ∼ 0.2) whisker position but provided moderate predictive power (R^2^ ∼ 0.4) for locomotion speed (Fig. 5f), indicating that whisker position is represented by a specialized population of neurons that are not driven simply by changes in brain state related to total body movement.

The neural representation of whisking in SC neurons likely depends on multiple brain areas (*36, 79*). The trigeminal nuclei and somatosensory cortex are the primary sources of whisker afference to the SC (*17*). Activity in these brain regions is strongly correlated to the phase of whisker motion (*22, 24, 80, 81*). In addition, the motor cortex and cerebellum likely send whisker efference to the SC (*12, 19*). Neural activity in these brain regions is correlated to the amplitude and midpoint of whisker motion (*27, 29–31, 82, 83*). Importantly, projections from all four of these brain regions converge in a lateral zone of the SC that closely matches our recording site (*18*). Therefore, the representation of whisker kinematics in SC neurons is likely built from the integration of afferent and efferent inputs. This model of convergence is further supported by our whisker trimming results, which caused a modest disruption to slow self-motion tuning and a more frequent change to phase tuning (Fig. 6). This is consistent with the sensory origin of phase tuning in the barrel cortex (*80*). The activity of most SC neurons was biased towards present or future kinematic features (Fig. 3), suggesting a correlation to reafferent and efferent information. In conclusion, these data indicate that the SC combines multiple signals to build a comprehensive representation of whisker position that reflects past, present and future states of self-motion. Such computations are critical for active sensing and computing the difference between the expected and real-world outcome of body movement (*44, 61, 63, 84*).

The SC is known to play an important role in driving orienting movements. Microstimulation of its deep layer neurons shifts the gaze of primates by activating neck and eye muscles (85). Therefore, the SC serves as a locus for coordinating distinct but related effector muscles. In our study, we discovered individual SC neurons that encoded both whisking and locomotion (Fig. 3), indicating that mouse SC may also coordinate complex movements through the control of complementary motor systems (*54*). Anatomical evidence supports this hypothesis, whereby SC neurons target brain regions involved in explorative or appetitive locomotion (*57, 77, 86*). Furthermore, the SC provides monosynaptic excitation to the nucleus that drives the whisking muscles (*35, 37, 40*). In line with this evidence, we discovered a subset of SC neurons that reliably spiked milliseconds before the start of whisker protraction, with greater spike rates preceding larger protractions (Fig. 4i).

Moreover, the activity of many SC neurons was predicted well by future whisker positions, revealing their potential role in movement control (Fig. 3e). Therefore, SC neurons have the requisite anatomy and physiology for coordinating whisking and locomotor strategies (*54*).

The gestalt of our data indicates that many SC neurons calculate an internal estimate of whisker position. A key feature of internal models is their ability to distinguish external from self-generated stimulation (*87*).

Consistent with this feature, we previously revealed a transient response in SC neurons that emerged during unexpected, externally generated touch (*49*). In the current study, we discovered that this external response is modulated by the position of the whisker upon touch. Importantly, the correlation between whisker position and SC firing rate was similar between the free-whisking and touch conditions. Therefore, the internal state established by self-motion directly modulates the strength of the external response. This modulation was linear and functionally could signal the egocentric distance of an object. Taken together, our data reveal that SC neurons incorporate self-motion information in real-time, constantly adjusting their output to represent the sensorimotor variables underlying active tactile perception.

### Limitations of the study

The main limitation of our study is that it lacks causal evidence delineating the contribution of afferent and efferent information to self-motion tuning. While it is clear from our trimming results that afferent information is an important component, we could not determine its absolute contribution to different kinematic features. To overcome this limitation, an approach is needed that reversibly deactivates the infraorbital sensory nerve without disturbing data collection. A post-hoc analysis comparing LNP model performance pre- and post-deactivation would reveal the absolute contribution of sensory reafference. Likewise, reversible deactivation of the facial motor nerve is necessary to clearly delineate the contribution of a motor command signal. Facial motor nerve deactivation would eliminate whisking-related reafference, attributing any remaining model performance to motor efference. Despite this limitation, our time-shifted models provide important insight into the role of SC in whisking. At surface level, neurons predicted by past kinematic features are likely to integrate sensory reafference, while neurons best predicted by future kinematics are potentially generating self-motion. Therefore, our work provides a critical first step towards untangling the precise sensorimotor contribution of individual SC neurons to active touch.

## Methods

### Experimental model and subject details

Mice of CD-1 background of both sexes between the ages of 9 and 15 weeks were used for all experiments. The Purdue Institutional Animal Care and Use Committee (IACUC, 1801001676A004) and the Laboratory Animal Program (LAP) approved all procedures. Mice were housed at room temperatures ranging between 68 and 79°F with humidity ranging between 40 and 60%. Mice were socially housed with five or less per cage and maintained in a reverse light-dark cycle (12:12□h). All experiments were conducted during the animal’s subjective night.

### Preparation for In-vivo electrophysiology

Each mouse was fitted with a custom designed aluminum headplate for head fixation. Animals were anesthetized with isoflurane (3 – 5%) while their body temperature and respiratory rate were monitored. To prevent eye dryness, artificial tears ointment was applied. The skin and fur on the skull were disinfected using 70% ethanol and betadine, and then incised with sterilized surgical instruments. Liquivet tissue adhesive and Metabond dental cement were applied to the skull and wound margins to secure the headplate. Buprenorphine was administered for pain relief as a post-operative analgesic.

One day before the recording session, mice were briefly (15-20 minutes) anesthetized to perform a craniotomy over the SC. A 1 mm diameter craniotomy was made (coordinates 4 mm posterior and 1.5 mm lateral from bregma) with a Robbins Instruments biopsy punch and sealed with Dowsil silicone gel and Body Double. The next day, mice were head-fixed on the free-spinning circular treadmill in the electrophysiology set-up. A three-shank, 128-channel Neuronexus probe was inserted at the site of craniotomy using a NewScale micromanipulator. The probe was lowered through the cortex at 75 µm/min, searching for light-driven activity. After passing through the cortex, denoted by a total loss of spiking, the return of visual responses indicated electrode penetrance into the superficial layer of the SC (∼1000 µm below the cortical surface). The probe was further descended into the intermediate and deep layers, where whisker deflections elicited spiking. The receptive field of recorded neurons was mapped by deflecting individual whiskers and to identify which whiskers elicited the greatest response. Whiskers that did not elicit detectable changes in neural activity were trimmed off to improve whisker tracking. Recordings were targeted at the C-row whiskers. If the electrode missed the target, it was removed, and re-positioned based on somatotopic coordinates. In most experiments, mice retained 3–5 whiskers across one or two rows; in a few experiments, only one or two whiskers were left intact.

### Mouse behavior

Two days after head plate implantation, mice were given the opportunity of run on a circular treadmill for 1 hour daily, until they fully habituated to the apparatus and were able to maintain a steady running pace. This facilitated volitional running and active whisking on the day of the electrophysiological recording. An opaque flag was placed over the eye and white noise was broadcasted to minimize visual and auditory cues, respectively. To measure the distance travelled by the mouse on the wheel, the circular treadmill was attached to a rotary encoder. A trial was initiated after the mouse ran 200 cm on the treadmill, and ended after the mouse ran an additional 200 cm. Whiskers were imaged during the trial window. In 20% of trials, chosen in a random order, a stepper motor moved the tactile surface into the active whisking field. At the end of the tactile stimulus trial, the surface returned to its location outside the whisking field.

### Spike Sorting

Spikes sorting was performed using the MATLAB package Kilosort2 and manually curated using Phy2 gui (https://github.com/cortex-lab/phy) (*88*). Spike clusters were considered single units based on their spike waveform, location on the electrode, and cross-correlogram of spike times. Single units were used for all analyses in the paper.

### Whisker Imaging & tracking

Whiskers were imaged using a high-speed (500 fps) infrared camera (photonfocus DR1) through a mirror angled at 45^0^ under IR illumination. Imaged videos were synchronized with neural spike data via external triggers using a National Instruments card and recorded on an Intan 512 controller. DeepLabCut was used to track whisker movement and curvature (*51*). Whisker curvature was calculated from the three distal labels (of four total) on each whisker using Menger curvature. About 150 frames from each recording session were labelled manually, spanning diverse whisker positions. The neural network was trained for at least 200k iterations, and the final labels were manually verified for accuracy.

### Extracting self-motion features

Whisker position was calculated as the angle between the frame’s vertical axis and the line joining a point on the mouse’s nose to a label on the whisker (see Suppl. Video 1 for reference). The chosen label was from a whisker that did not make contact with the tactile surface and it was the same between the intact and trimming conditions. Whisker envelope was generated by interpolating peaks and troughs of the whisker position using Akamai spline interpolation (see Figure 1b). Whisker midpoint was calculated as the median of the upper and lower bounds of the envelope, while amplitude was the half width of the envelope.

Whisker phase was obtained by bandpass filtering the whisker position trace between 5 and 50 Hz (bandpass(), MATLAB) followed by calculating the Hilbert transform (hilbert(), MATLAB) and obtaining the angle from the complex value (angle(), MATLAB). A phase angle of 0° is the most protracted angle, while +/-π was the most retracted angle.

### Tuning curves

To create tuning curves for slow features like locomotion speed, whisker midpoint and amplitude, we segmented neuronal spike rates into 50ms time intervals. Each feature’s total range was divided into 20 equal bins. For each feature bin, we calculated the mean and standard error (SEM) of the spike rate. The slopes of the tuning curves were generated by fitting the firing rates and feature values in 50ms bins using a linear regression model (fitlm(), MATLAB). To find sharp transitions in whisker midpoint, findchangepts() function in MATLAB was used.

To obtain a neuron’s phase tuning, we generated spike triggered averages of whisker phase. The spike counts for each neuron were binned into 30° non-overlapping phase bins spanning from – π to +π, determined by the whisker phase angle at the time of each spike. To obtain neuronal firing rates as a function of whisker phase, spike counts in each phase bin were divided by the time spent at each angle. To test for significant phase tuning, we compared each neuron’s firing rate distribution to a uniform distribution using chi-squared goodness of fit. The phase preference of each neuron was the circular mean of its phase tuning curve (circ_mean(), CircStat toolbox, MATLAB). The phase modulation depth of each neuron was calculated using the maximum, minimum and mean firing rates of the phase tuning curve (*22*).

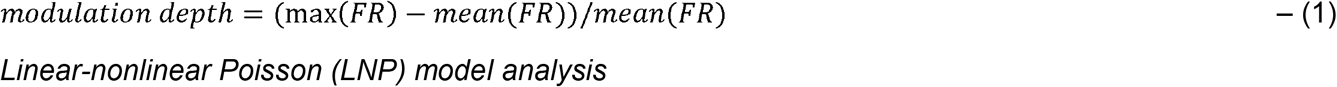

To quantify the dependence of spiking on a feature, or a combination of features (locomotion speed, whisker midpoint, whisker amplitude, and whisker frequency) we employed a linear-nonlinear Poisson spiking GLM model (LNP), previously established in the field (*53*). This approach is agnostic to tuning curve shape and robust to the interdependence of encoded variables. Considering the significant correlation between self-motion features observed in our study, this LNP approach is highly advantageous over traditional tuning curves (Suppl. Fig 3a).

The LNP model estimates the spike rate (N^t^) of individual neurons at each time bin ‘t’ by taking an exponential sum of the weighted feature values. In the below equation, ‘f’ represents the features (Locomotion speed, Midpoint, Amplitude, Frequency), X_f_^t^ is a feature vector at time ‘t’, W_f_ is the learned weight that converts feature value to firing rate, and dt is the duration of time bin (50ms). We binned neuron spike rates and self-motion features into 50ms bins.

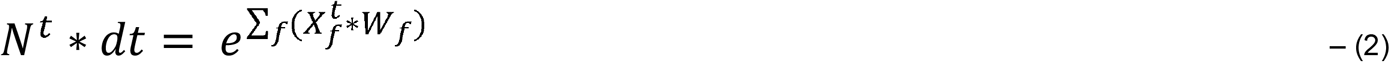

X_f_ is a binned feature occupancy vector. Each feature is binned into 20 bins, that span the feature’s entire range. At each time point t, all bins are set to 0 except for the one that corresponds to the feature value at that time, which is set to 1. To determine the weight vector W_f_ for each neuron, we maximized the Poisson log-likelihood of the observed spike train. The model performance for each neuron was assessed as the log-likelihood (LLH) on held out data. We performed 10-fold cross-validation.

To identify which subset of features best explains spiking in each neuron, we performed a forward search approach (Suppl. Fig 3c,f). This approach tested models with varying numbers of features: four 1-feature models (L,M,A,F), six 2-feature models (LM, LA, LF, MA, MF, AF), four 3-feature models (LAM, LAF, MAF, LMF), and one full model (LMAF). The process began by selecting the single-feature model with the highest performance. This model’s performance was then compared against all two-feature models that included the chosen single feature. If the best-performing two-feature model significantly outperformed the single-feature model, we then compared it to the three-feature models, and so forth (Suppl. Fig 3e,f). The chosen model was the one with the fewest features that did not show a significant improvement in prediction when additional features were added. If a neuron’s best model didn’t significantly outperform the mean firing rate model, it was labeled as unclassified. We determined significance using a one-sided signed rank test with an α of 0.05. For more details on the model, refer to https://github.com/GiocomoLab/ln-model-of-mec-neurons.

### Relative contribution of single features

We identified the contribution of a given feature by finding the difference in model performance (LLH increase) between the selected model and the model that contains all the features in the selected model except the given feature. For example, if a neuron’s selected model is LMA, then contribution of feature M is contrib(M) = LLH(LMA) – LLH(LA). Contributions were calculated for a given feature even when the given feature was not contained within the selected model for a neuron. For example, if a neuron’s selected model is LA, then the contribution of M is contrib(M) = LLH(LMA) – LLH(LA). Contributions of features that are not encoded in a neuron are usually very small, consistent with model selection procedure. In case contribution of a given feature was negative (which could happen when the feature was not encoded by the neuron), then the negative values were reset to 0. After computing contributions of all 3 features, we normalized the contributions by making their sum equal 1.

### Time shifted models

For each neuron, we fit 12 time-shifted LNP models. We binned the data into 25ms, to accommodate for time shifts lower than 50ms. The feature values were shifted by {-200, -150, -100, -75, -50, -25, 0, 25, 50, 75, 100, 150, 200} ms relative to the spike train. We plotted the log-likelihood (LLH) of each model against its time shift to create a temporal curve. Temporal bias was calculated as the difference between the area under the curve (AUC) for positive and negative time shifts, and then dividing this difference by the total area. The time shift with the highest LLH was identified as the preferred time shift.

### Self-motion decoder model

To determine if self-motion encoding in SC neurons support downstream decodability, we implemented a ridge regression model (Fig. 5a, sklearn.linear_model.Ridge, Python). We decoded whisker position with a temporal resolution of 15ms, matching the resolution of our binned spike rates. For decoding, we utilized a 165ms time window, comprising 11 bins (5 preceding, 1 concurrent and 5 following bins) of spike rate data. To fit a model, we conducted 10-fold cross-validation, and decoding performance was tested on the 20% of held-out data.

When decoding with a reduced number of neurons (from population total), we randomly selected self-motion encoding neurons, and the randomization was repeated 10 times to accurately gauge decoding performance. For more details of the code, refer the python version at https://github.com/sumachinta/body-position_decoding_model/tree/main.

### Simulated neuronal firing rates for self-motion decoding

Neuronal firing rates were simulated based on self-motion tuning curves. Firing rates were simulated in 15ms time bins. The firing rate in a time bin t was generated by picking a random number from a Poisson distribution specified by the rate parameter N^t^ calculated from equation 2 (poissrnd(), MATLAB). If a neuron did not encode a specific feature ‘f’, the weight ‘Wf’ was set to 0.

To simulate neuron firing rates tuned to whisker phase, we divided the phase range into 12 bins and created a vector for each time bin ‘t’. In this vector, all bins were set to zeros except for the one corresponding to the current phase at time ‘t’, which is set to 1. Multiplying this vector with each neuron’s phase tuning curve gave us the rate parameter. The firing rate for each time bin was then determined using a random number from a Poisson distribution based on this rate parameter. The feature values to simulate spiking were taken from a randomly chosen recording session.

### Sensory reafference

To examine how reafference affects the coding of self-motion in SC neurons, we trimmed off the whiskers for the final ∼25 trials of the recording session, sparing a portion of the whisker for tracking movement. To compare the change in self-motion tuning to slow kinematic variables, we fit a separate LNP model during the whisker trim period. We compared the model performances before and after whisker trimming by looking for significant differences in LLH over 10 folds of training (t-test, 10-fold cross-validation). Neurons were categorized as modified if they exhibited a significant change in LLH, or unchanged if they did not. Neurons that had a selected LNP model before whisker trimming and no selected model after trim were categorized as lost units, while gain units had the opposite trend.

To measure the change in phase tuning, we checked if there was significant correlation between the tuning curves before and after whisker trimming (using Pearson correlation coefficient corr(), MATLAB, with α = 0.05). Neurons that had a significant correlation between pre- and post-trim conditions were labeled as ‘no change’ units. Those without a significant correlation were identified as ‘modified’ units. Neurons that were phase-tuned before whisker trimming (see ‘phase tuning curve’ section of methods) but not classified post-trimming were categorized as ‘lost’ units. Conversely, neurons that were not phase-tuned pre-trimming but became tuned post-trimming were classified as ‘gain’ units. Neuron’s spike waveforms were compared pre- and post-trimming to ensure that the units were stable across pre- and post-trimming conditions. Mean firing rates of neurons during pre and post trim condition were controlled for whisker position and locomotion speed. To match the whisker position and locomotion speed coverage, we generate two-dimensional occupancy matrices where each bin is a combination of a position and speed bin. Finally, we matched position and speed coverage across two conditions by down-sampling data points from either condition so that occupancy time was matched for each bin.

### Tactile response

To investigate whether self-motion affects tactile responses in the SC, we studied how these responses change as a function of whisker position and locomotion speed during the tactile response window. We determined a fixed response window for all neurons in each recording session, based on population mean firing rates. Within this window, we calculated the average firing rate of neurons, as well as the average whisker position and running speed. We then used the Pearson correlation coefficient (p < 0.05, corr(), MATLAB) to assess whether the firing rate was correlated to either whisker position or locomotion speed. Previous work in SC has shown that tactile responses habituate with stimulus repetition (*49*). To account for this, we excluded neurons that displayed a stronger correlation with stimulus repetitions than self-motion (Suppl. Fig. 6).

## Acknowledgements

The authors would like to acknowledge the members of the Pluta lab, James Dooley, Kate Hong and Krishna Jayant for helpful discussions and feedback on the manuscript. They would also like to thank Shreya Beri for assisting with whisker tracking.

## Funding

Purdue Institute for Integrative Neuroscience

The Whitehall Foundation

Showalter Trust

## Author Contributions

Conceptualization: SC, SRP

Methodology: SC, SRP

Investigation: SC

Visualization: SC, SRP

Supervision: SRP

Writing: SC, SRP

