## Supplemental figures for "Internal monitoring of whisking and locomotion in the superior colliculus"

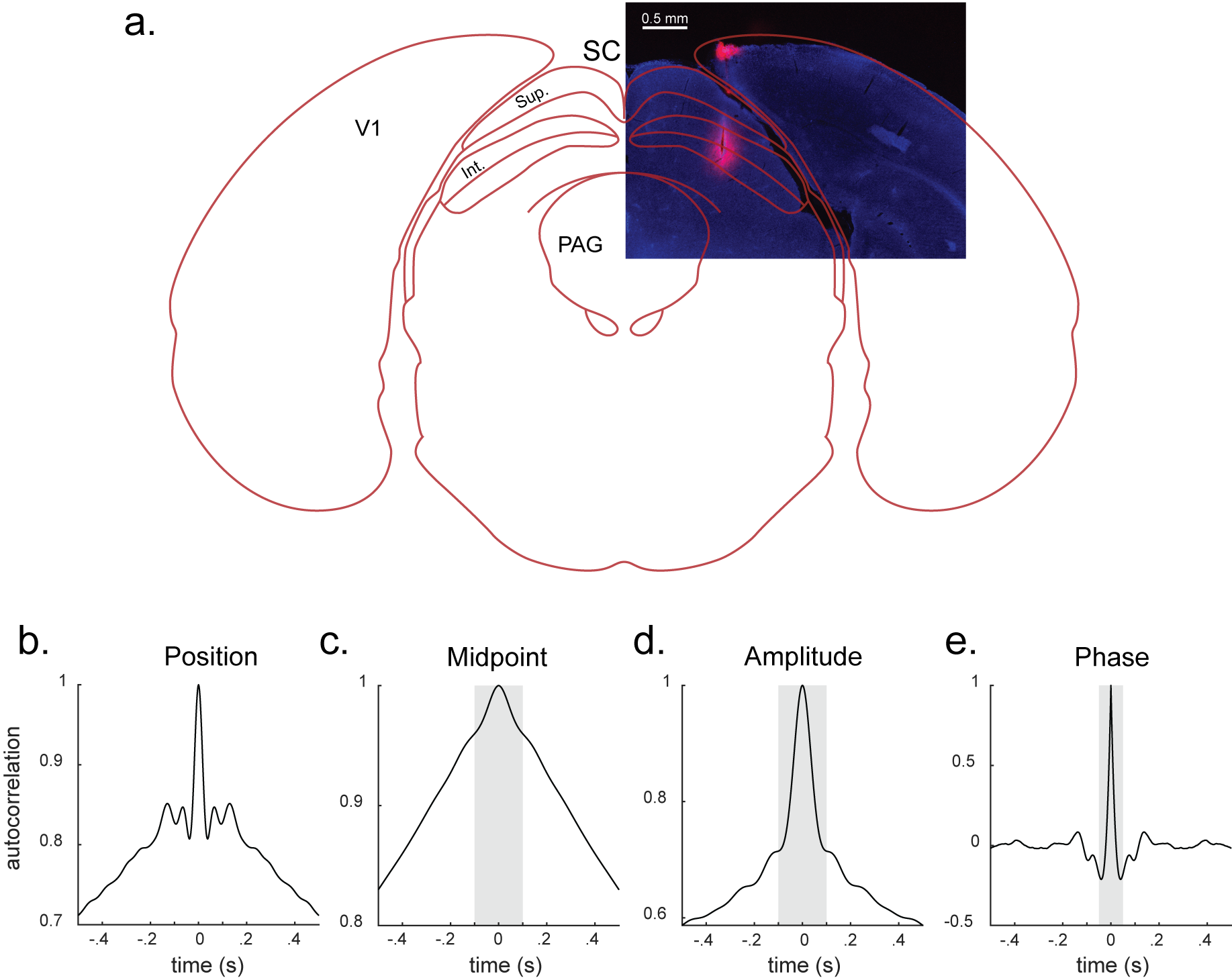

**Supplemental Figure 1. Electrode penetration and self-motion feature timing**. **A**) Dye labelling of the electrode shank at the recording site in the intermediate and deep layers of lateral SC overlaid with the outline of the coronal section taken from mouse brain atlas based on the stereotaxic coordinates of the recording site. **B**) Autocorrelation of whisker position reveals the fast and slow components. **C, D, and E**) Autocorrelation of whisker midpoint, amplitude, and phase. Midpoint and Amplitude vary more slowly than phase.

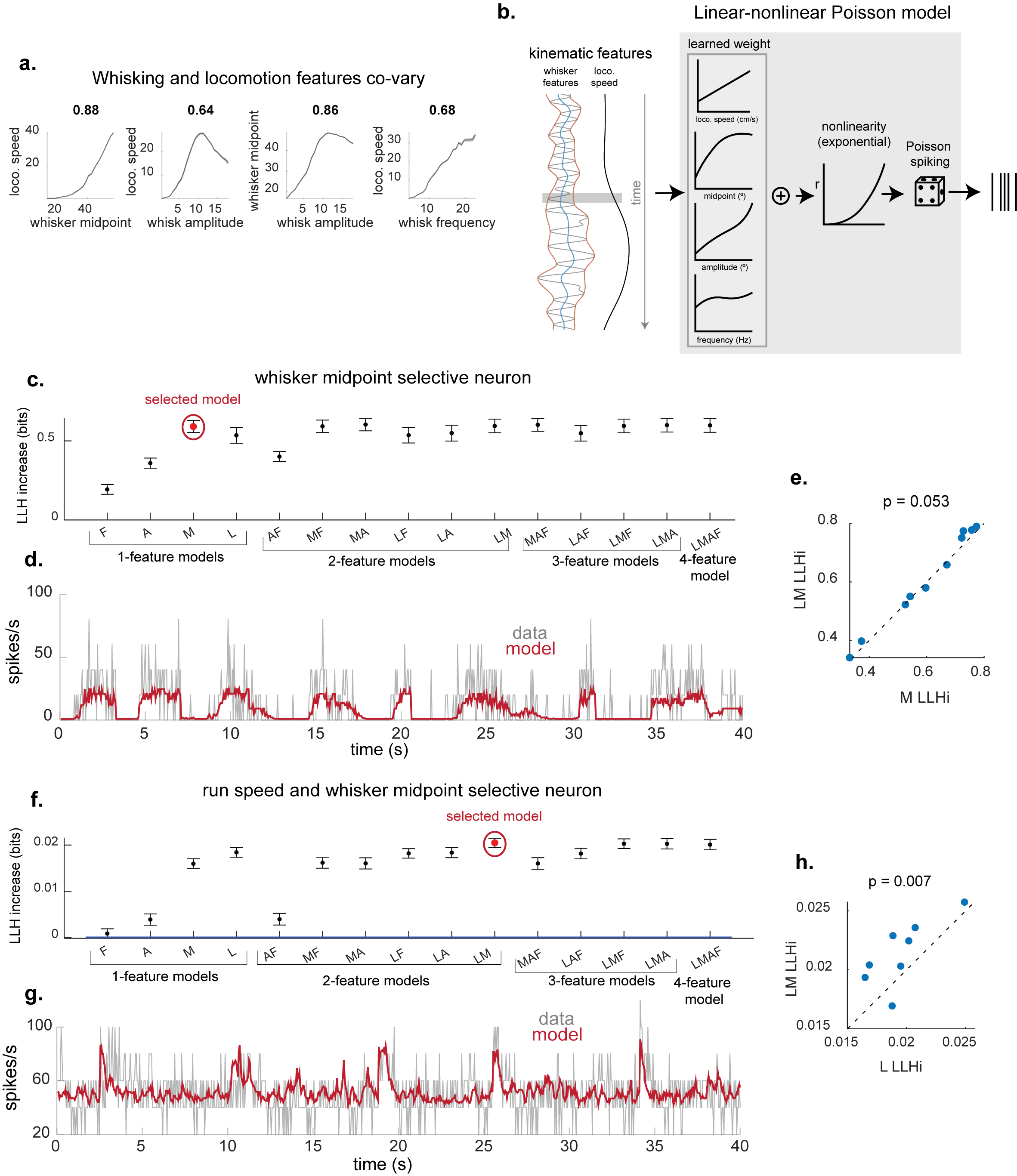

**Supplemental Figure 2. An LNP based GLM to identify individual neuron selectivity to self-motion features**. **A**) Mean curves depicting the co-variation of locomotion speed and whisking dynamics, with Pearson correlation coefficients indicating the strength of association (1 mouse). **B**) Schematic of a LNP framework model workflow, where kinematic features are weighted, summed, and passed through an exponential non-linearity to produce a Poisson-distributed spike estimate. C) Example midpoint selective neuron and its spike prediction performance across single and multi-feature models. The forward search approach identifies the minimum feature model whose performance is significantly better than any simpler model. The selected model for this neuron is marked in red. **D**) Overlay of the neuron’s actual firing rate (in grey) against the firing rate predicted by the selected model (in red), demonstrating the model's fidelity in capturing the neuron's response pattern. **E**) Scatter plot comparing the performance of the 'M' model against the 'L+M' model. A p-value of 0.053 suggests no significant benefit from including 'L', favoring the simpler 'M' model (right-tailed signed-rank test). **F, G, and H**) same as C, D, and E for a 2-feature selective neuron.

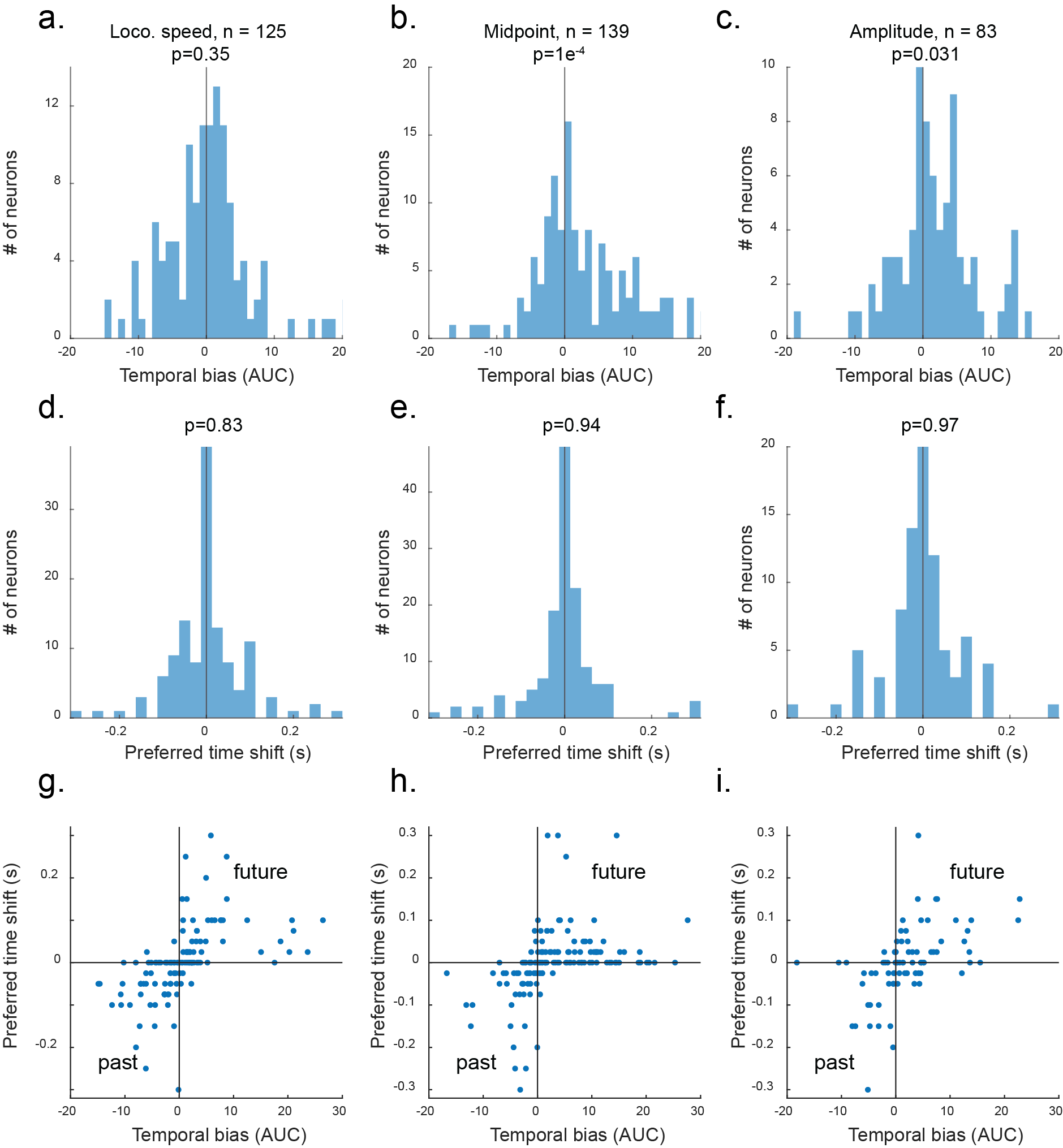

**Supplemental Figure 3. Temporal preference of neurons according to kinematic feature preference**. **A, B, and C**) Distribution of temporal biases for neurons whose best feature is locomotion speed, or midpoint, or amplitude respectively (125 neurons for locomotion speed, 139 neurons for midpoint, and 83 neurons for amplitude respectively). **D, E, and F**) Distribution of preferred time shifts for neurons in A, B, and C respectively. **G, H, and I**) Scatter plot of Preferred time shift and temporal bias for neurons in A, B, and C respectively.

­

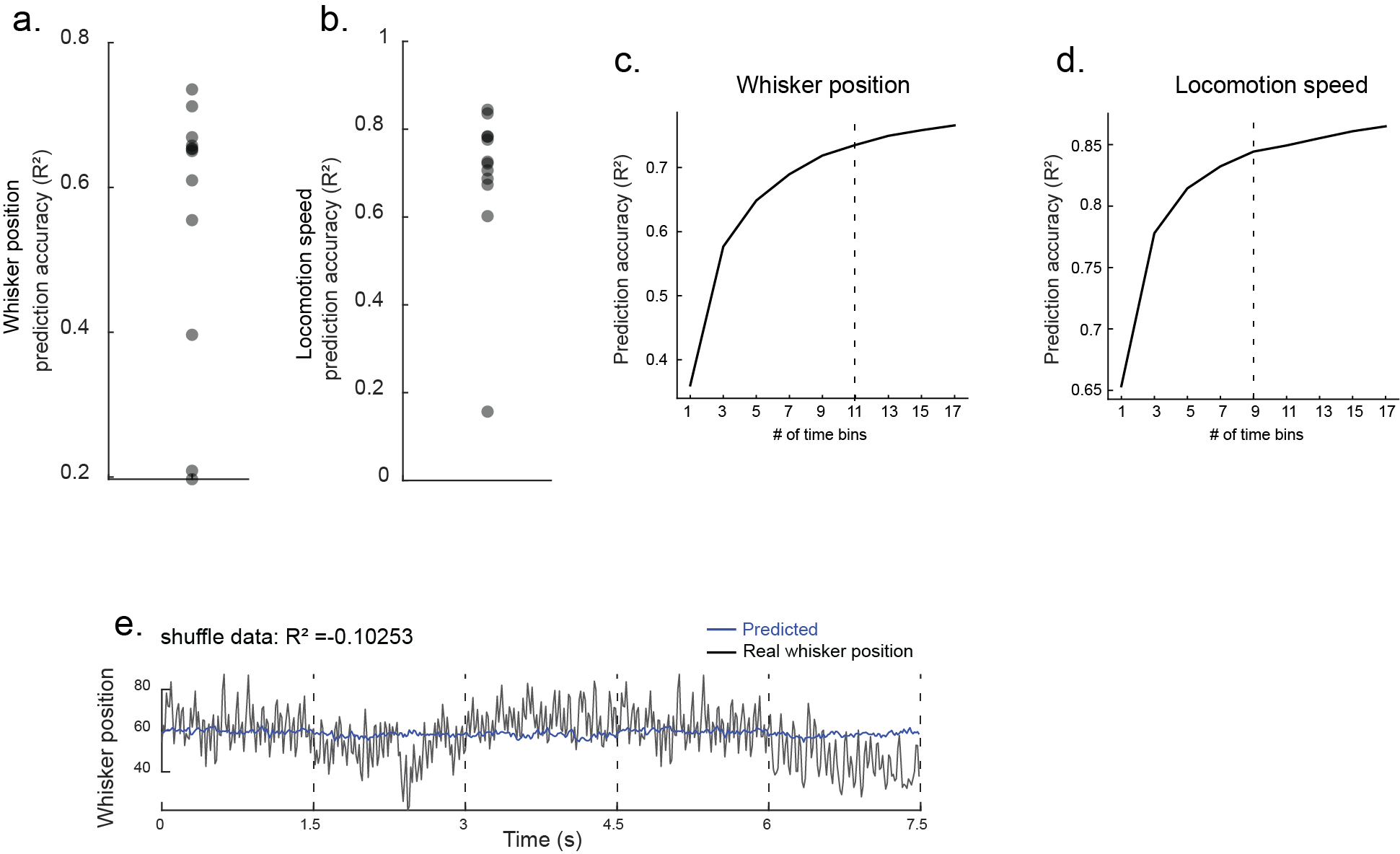

**Supplemental Figure 4. Decoding self-motion**. **A, B**) Distribution of whisker position and locomotion speed decoding accuracies for 12 individual mice. One mouse with the very low decoding accuracy for locomotion speed has only 1 locomotion speed tuned neuron. **C**) Whisker position decoding accuracy for 1 example mouse with increasing number of time bins (15 ms duration) in the self-motion decoder. **D**) Locomotion speed decoding accuracy for 1 example mouse with increasing number of time bins (100 ms) in the self-motion decoder. **E**) Whisker position decoding with simulated neurons with shuffled spikes.

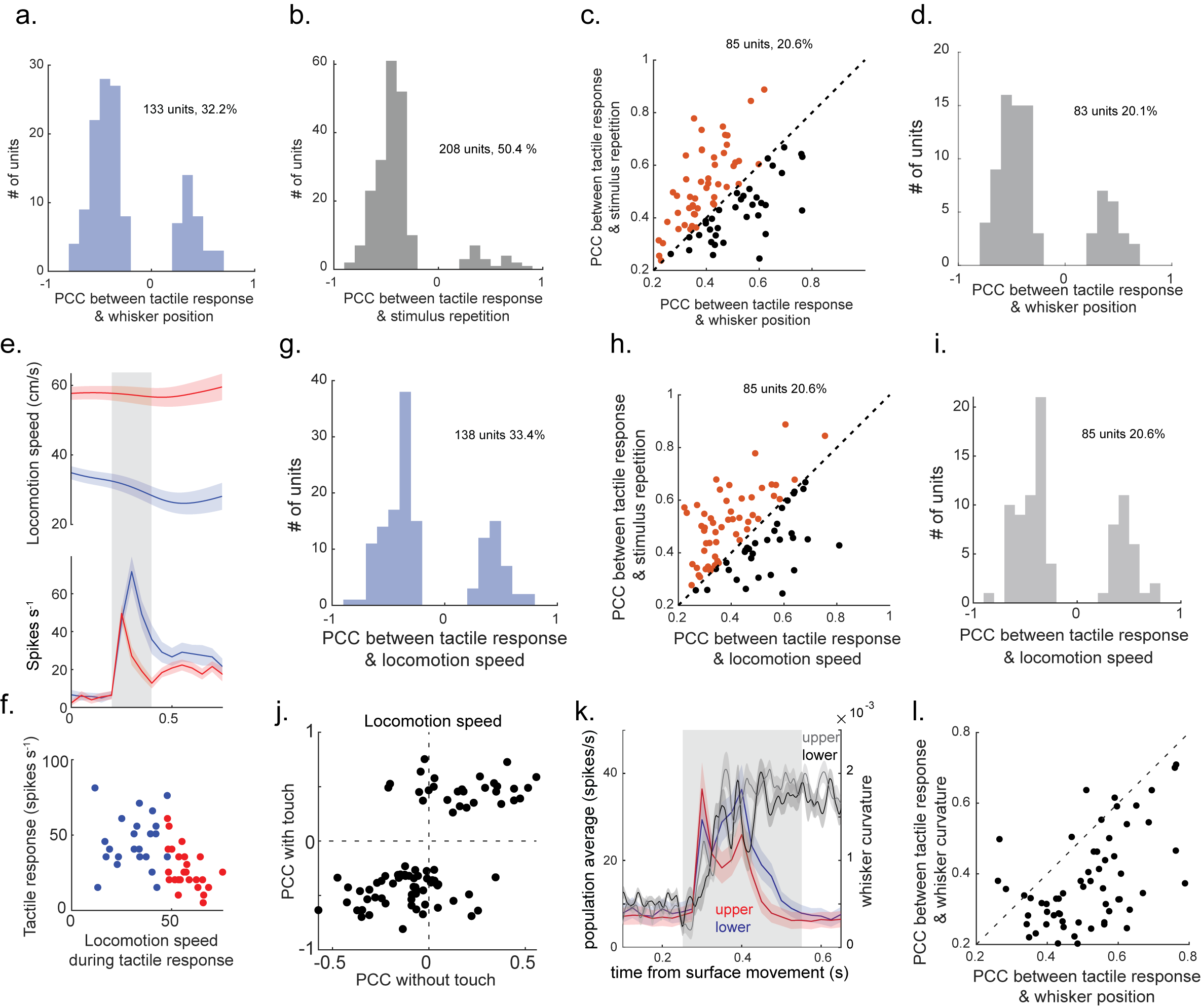

**Supplemental Figure 5. Response habituation and whisker curvature cannot explain dependency of tactile response on whisker position.** **A**) Distribution of Pearson correlation coefficients (PCCs) for neurons with a significant relationship between tactile response and whisker position (12 mice, 133 neurons, p < 0.05). **B**) Distribution of PCCs for neurons with a significant relationship between tactile response and stimulus repetition (12 mice, 208 neurons). **C**) Scatter plot comparing PCCs for the neurons that overlap between panels **A** and **B**. Neurons whose PCC with stimulus repetition is higher than whisker position PCC (in orange) are marked as repetition selective units and removed from further analysis. **D**) Distribution of Pearson correlation coefficients (PCCs) for neurons with a significant relationship between tactile response and whisker position (after removing stimulus repetition units, 12 mice, 83 units). **E**) An example neuron with strong correlation between tactile response magnitude and locomotion speed during the response window. Locomotion speed (top) and firing rate (bottom) during surface engagement. The tactile response window is marked in grey. In red are top 50% locomotion speed trials and in blue are bottom 50% locomotion speed trials. **F**) Scatter plot of mean spike rate vs mean locomotion speed during the tactile response period. **G**) Distribution of PCCs between spike rate and locomotion speed for neurons significantly correlated to locomotion speed in the tactile window. **H**) Scatter plot of the PCCs for locomotion speed and stimulus repetition for neurons that are significantly correlated to both locomotion speed and stimulus repetition. Neurons whose PCC with stimulus repetition is higher than locomotion PCC (in orange) are marked as stimulus repetition selective units and removed from further analysis. **I**) Distribution of PCCs for locomotion speed (after removing stimulus repetition selective units, 12 mice, 85 neurons). **J**) PCCs of locomotion speed during tactile response and during free whisking (12 mice, 85 neurons). **K**) Whisker curvature (black/gray) and population-averaged spike rate (blue/red) during the tactile response window for an example mouse. Red is the average of top 50% whisker curvature trials and blue is the average of bottom 50% whisker curvature trials. **L**) Scatter plot of PCCs for whisker position and whisker curvature. Trial-to-trial variation is more strongly correlated to whisker position.
